## Supplementary figures and images for "Closed loop motor-sensory dynamics in human vision"

### S Figure S1

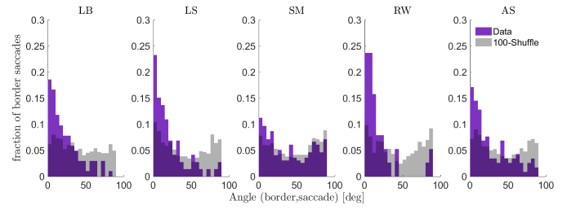

### S Figure S2

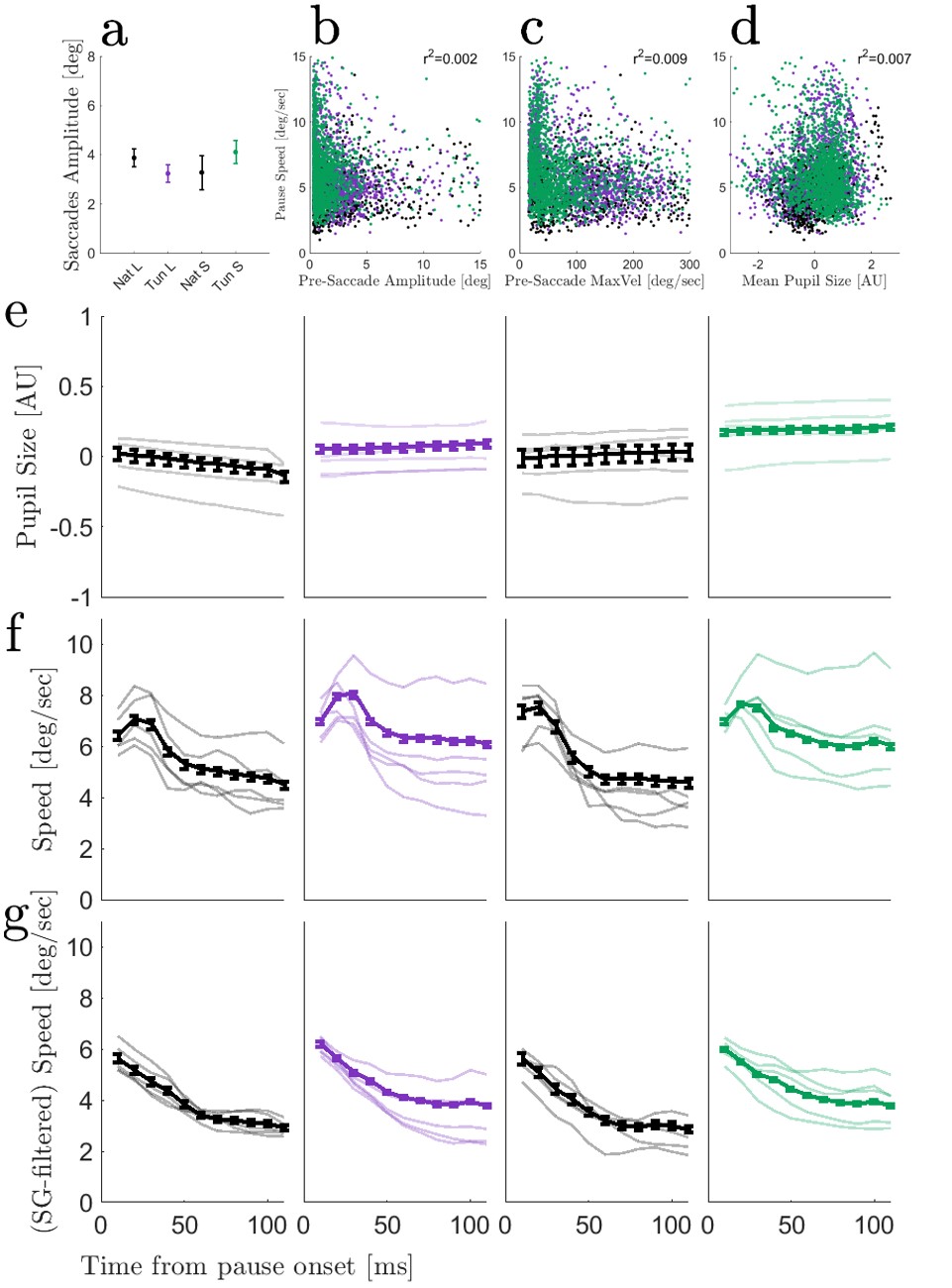

### S Figure S3

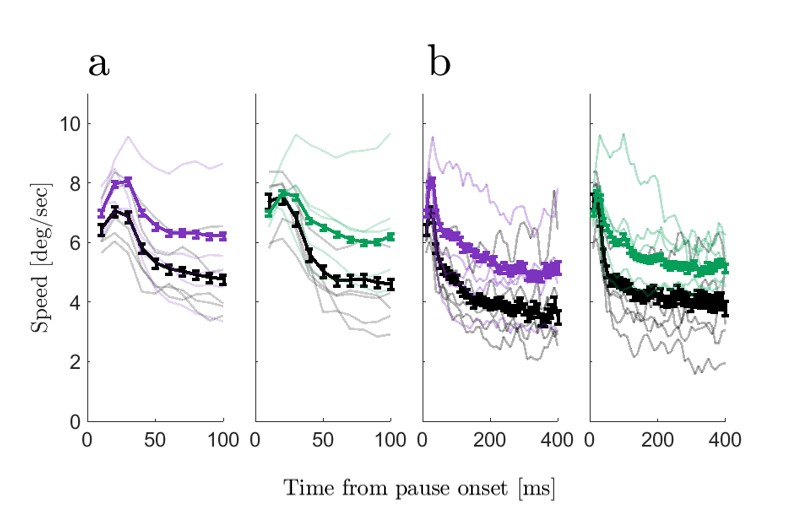

### S Table S1

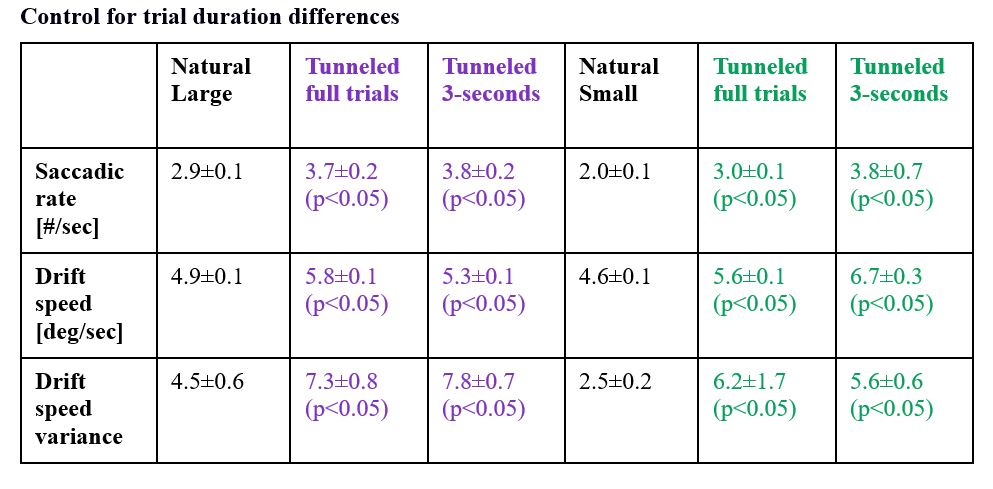
